# DLA-Ranker: Evaluating protein docking conformations with many locally oriented cubes

**DOI:** 10.1101/2021.10.26.465898

**Authors:** Yasser Mohseni Behbahani, Élodie Laine, Alessandra Carbone

## Abstract

Proteins ensure their biological functions by interacting with each other, and with other molecules. Determining the relative position and orientation of protein partners in a complex remains challenging. Here, we address the problem of ranking candidate complex conformations toward identifying near-native conformations. We propose a deep learning approach relying on a local representation of the protein interface with an explicit account of its geometry. We show that the method is able to recognise certain pattern distributions in specific locations of the interface. We compare and combine it with a physics-based scoring function and a statistical pair potential.

## 1 Introduction

Protein-protein interactions play a central role in virtually all biological processes. Reliably predicting who interacts with whom in the cell and in what manner would have tremendous implications for bioengineering and medicine. Hence, a lot of effort has been put into the development of methods for docking proteins against each other and identifying the most probable 3D arrangements they form *in vivo*. While highly efficient algorithms can exhaustively sample the space of candidate conformations (Ritchie and Venkatraman, 2010), correctly evaluating and ranking these conformations remains challenging. In this work, we investigate the possibility of discriminating near-native complex conformations from decoys by exploiting local 3D-geometrical and physico-chemical environments around interfacial residues. Our motivation is that the number of known protein-protein complex structures is fairly limited. Breaking down these structures into interfacial residue-centred local environments allows training on a much larger set of labelled data.

We propose Deep Local Analysis (DLA)-Ranker, a deep learning-based approach predicting whether a candidate complex conformation is acceptable or not. To do so, it applies 3D convolutions to a set of locally oriented residue-centred cubes representing the interface between the proteins (**Fig. 1**). We orient the cubes by defining local frames based on the common chemical scaffold of amino acid residues in proteins (Pagès *et al*., 2019). This representation guarantees that the neural network output is invariant to the global orientation of the input conformation while fully accounting for the relative orientation of the central residue with respect to its neighbours.

**Figure 1:**
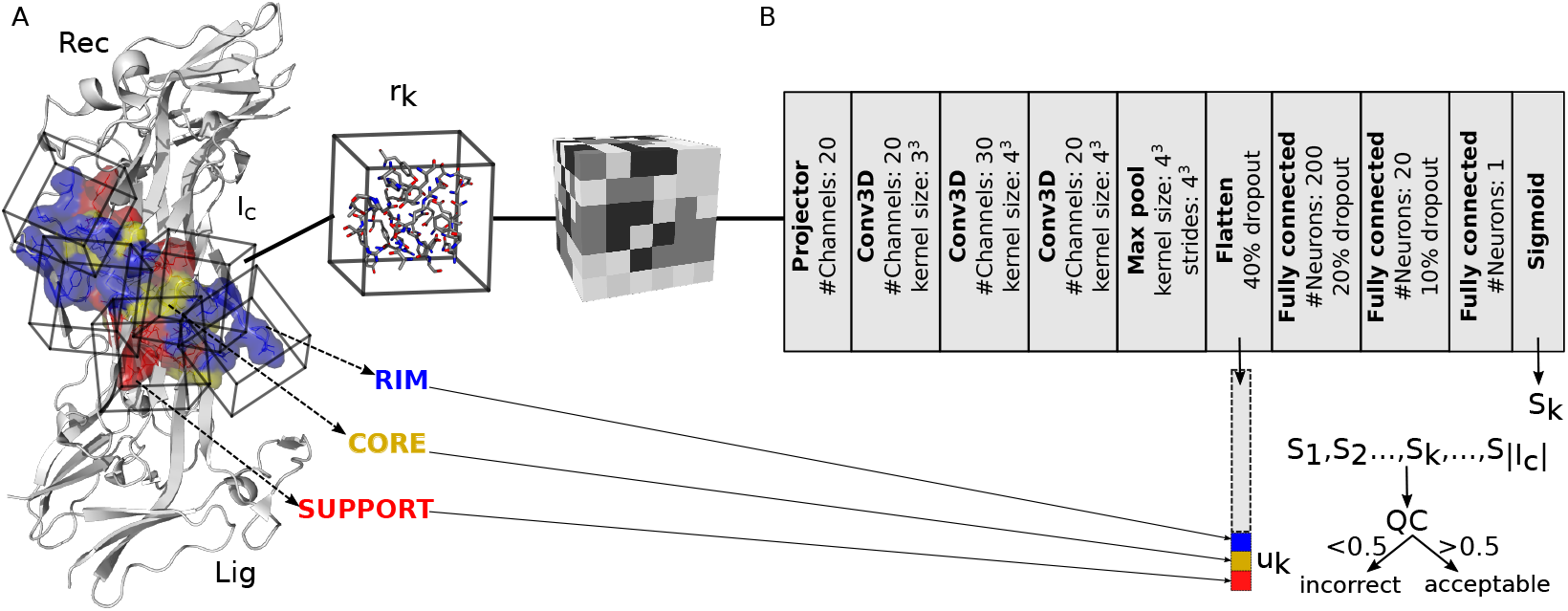
A) Representation of the interface as an ensemble of cubes. Each cube is labelled S, C, R, depending on the position of the central residue within the interface. B) Architecture of DLA-Ranker neural network.

## 2 Related works

A few of 3D convolutional neural network-based approaches have been developed to evaluate docking conformations (Renaud *et al*., 2021; Wang *et al*., 2020). Contrary to our approach, they intend to represent the whole interface as one single voxelized 3D grid without defining any local frame. Since this representation is sensitive to the orientation of the candidate conformation, and standard 3D convolutional filters are not rotationally invariant nor equivariant, the output may change upon rotation of the input in an uncontrolled fashion. To limit this effect, (Renaud *et al*., 2021) performed rotational data augmentation. Some other works have proposed alternative representations such as graphs (Wang *et al*., 2021; Cao and Shen, 2020) or point clouds (Eismann *et al*., 2021). By contrast to our local-based approach, they adopt a global perspective by assessing the quality of the interface (Wang *et al*., 2021) or even the complex (Cao and Shen, 2020; Eismann *et al*., 2021) as a whole. Representing the input conformation as a graph (Cao and Shen, 2020; Wang *et al*., 2021) renders the prediction rotationally invariant, but the information of the relative orientations of the atoms in the structure is lost. In the point cloud-based method (Eismann *et al*., 2021), the authors eliminate the need for rotational data augmentation by using tensor field network layers where the convolutional filters are decomposed into series of spherical harmonics.

## 3 Method

DLA-Ranker takes as input a cubic volumetric map centred and oriented on a given interfacial residue (**Fig. 1a**). Any residue displaying a change in solvent accessibility upon complex formation is considered as part of the interface. We used NACCESS (Hubbard and NACCESS, 1993) with a probe radius of 1.4 Å to compute residue solvent accessibility. To build the map, we adapted the method proposed in (Pagès *et al*., 2019). The atomic coordinates of the input conformation are first transformed to a density function. The density ***d*** at a point 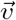 is computed as

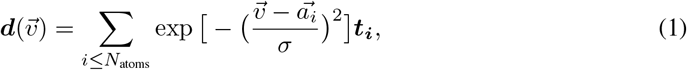

where 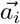 is the position of the *ith* atom, *σ* is the width of the Gaussian kernel and is set to 1Å, and ***t***_***i***_ is a vector of dimension 169 encoding some characteristics of the protein atoms. Namely, the first 167 dimensions correspond to the atom types that can be found in amino acids (without the hydrogens), and the 2 other dimensions correspond to the two partners, the receptor and the ligand.

Then, the density is projected on a 3D grid comprising 24×24×24 voxels of side 0.8Å. For the *nth* residue, the 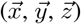 directions and the origin of the map are defined by the position of the atom N_*n*_, and the directions of C_*n-*1_ and C*α*_*n*_ with respect to N_*n*_ (Pagès *et al*., 2019). Thanks to this local frame definition, the map not only is invariant to the candidate conformation initial orientation but also provides information about the atoms and residues relative orientations.

The network architecture comprises three 3D convolutional layers (**Fig. 1b**). We trained it to perform the classification task of predicting whether a given interfacial residue is part of an acceptable (or better) docking conformation or an incorrect one, according to CAPRI criteria (Lensink *et al*., 2017). Depending on the location of the residue at the interface, we expect very different geometrical and physico-chemical environments. For instance, the map computed for a residue deeply buried in the interface will be much more dense than that computed for a partially solvent-exposed residue at the rim. This motivated us to explicitly give some information to the network about the location of the input residue. To do so, we classified the interfacial residues in three substructures, the support, the core and the rim (**Fig. 1a**), as defined in (Levy, 2010). We one-hot encode the input residue class in a vector *u* and append it to the embedding (or fingerprint) derived from the convolutional layers (**Fig. 1b**). The support-core-rim classification previously proved useful for the prediction and analysis of protein-protein and protein-DNA interfaces (Laine and Carbone, 2015; Raucci *et al*., 2018; Corsi *et al*., 2020).

We compiled our train, validation and test sets from two complete cross-docking experiments performed on about 400 proteins (Dequeker *et al*., 2019; Lopes *et al*., 2013) using the rigid-body coarse-grained docking tool MAXDo (Sacquin-Mora *et al*., 2008). We efficiently screened 27 millions docking conformations with INTBuilder (Dequeker *et al*., 2017) and the rigidRMSD library (Popov and Grudinin, 2014), and we systematically evaluated their quality with respect to the experimentally resolved complex structures available in the Protein Data Bank (Berman *et al*., 2002). To build our training set, we extracted 3 902 acceptable or better conformations and 6 038 incorrect conformations coming from 312 protein pairs (**Fig. S1**). The unbound forms of the proteins or their close homologs (*≥*70% sequence identity) were used for docking in about half of the pairs. We trained 5 models over 20 epochs following a 5-fold cross-validation procedure (**Fig. S2**). We minimised the binary cross entropy loss function using the Adam optimiser with a learning rate of 0.001 in TensorFlow (Abadi *et al*., 2015). We explored about 10 different architectures, by varying the number of convolutional layers, the number of neurons in the fully connected layers, and the dropouts. The calculations were performed on workstations with GPU: NVIDIA GeForce RTX 3090 (24 GB RAM) and CPU: AMD Ryzen 9 5950X. The generation of the interface cubes for a complex takes 1.46s on average, and their evaluation 0.23s.

For each input interfacial residue, the DLA-Ranker network outputs a score comprised between 0 and 1. If the score is higher than 0.5, we consider that the network classified the residue as belonging to a near-native (acceptable or better) conformation, incorrect otherwise. To evaluate an entire candidate conformation, DLA-Ranker averages the individual residue scores over the interface. Hence, the predicted quality *Q* of conformation *C* is expressed as

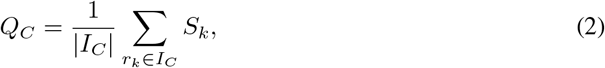

where *I*_*C*_ is the ensemble of interfacial residues and *S*_*k*_ is the score predicted by the network for the input 3D grid centred on the residue *r*_*k*_.

## 4 Results

We assessed the ability of DLA-Ranker to correctly rank candidate conformations on a test set of 20 protein pairs that were not seen during the training (**Table S1**). For each complex, we selected the 1 000 best-scored conformations according to the physics-based scoring function implemented in MAXDo (very similar to ATTRACT scoring function (Zacharias, 2003)) and evaluated them using DLA-Ranker. For most of the pairs, DLA-Ranker assigned high *Q* scores to the near-native conformations and is able to discriminate them from the incorrect ones (**Fig. S3**). We further ranked the conformations using a consensus of the 5 trained DLA-Ranker models. To do so, we first ordered the conformations according to the *Q* scores computed from each trained model. Then, we discretised the ranks into 7 bins, namely top1, top5, top10, top50, top100, top200, top1000, and lexicographically ordered these labels. DLA-Ranker put a near-native conformation in the first place for two thirds of the protein pairs (**Fig. 2A**). It achieved better performance than MAXDo in 11 cases. A particularly difficult case for both MAXDo and DLA-Ranker is the pair 1rkc_A:1ydi_A. Combining DLA-Ranker with the pair potential CIPS (Nadalin and Carbone, 2018) allowed enriching the top 200 subset for that pair in near-native conformations, and overall improved the results (**Fig. 2B**).

**Figure 2:**
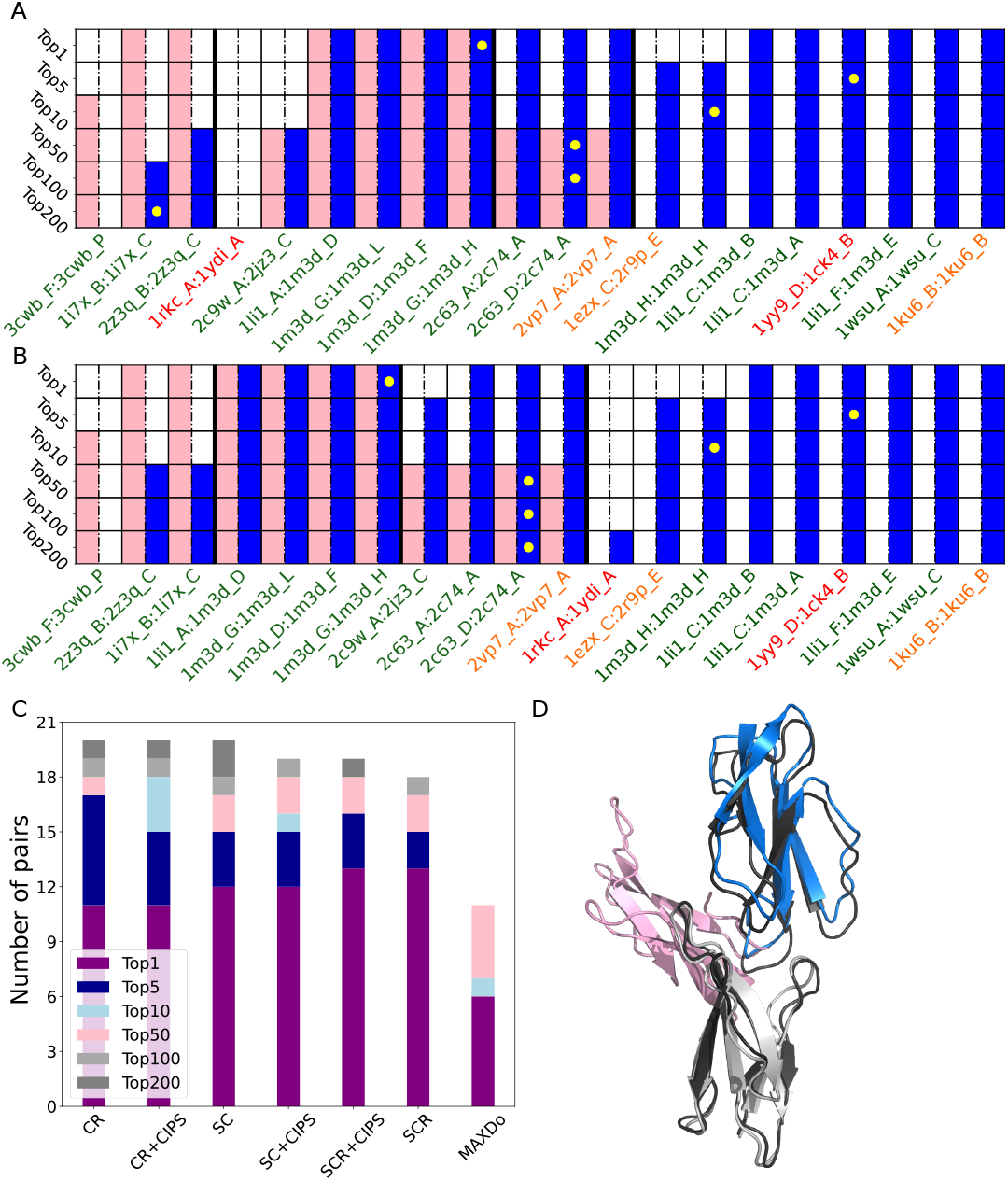
**A-B**. Ranking results per protein pair. For each pair, we report whether some near-native conformations were found in the top 1, 5, …, 200 out of a total of 1 000 conformations generated and selected by MAXDo. A coloured cell indicates the presence of at least one acceptable conformation in the corresponding topX. The pink color corresponds to MAXDo while the blue color corresponds to DLA-Ranker (A) or DLA-Ranker combined with CIPS (B). For each topX, the yellow dot indicates the pair with the highest enrichment factor. The PDB ids are coloured according to the magnitude of the conformational change between the docked forms and the bound forms. Green: none or small. Orange: medium. Red: large. **C**. Comparison between different methods. The SCR, SC and CR DLA-Ranker models were trained and tested on all interfacial residues, only those in the support and core, or only those in the core and rim, respectively. **D**. Best-ranked candidate conformations for the 1ku6_B homodimer. The reference complex structure is in black, the docked receptor in grey, the ligand conformation selected by MAXDo in pink and that selected by DLA-Ranker in blue.

Overall, DLA-Ranker performance seem to be independent from the extent of conformational change between the docked protein forms and the bound forms (**Fig. 2A-B**, label colors). For instance, one of the cases where it performs very well, the 1ku6_B homodimer, displays a substantial rearrangement (**Fig. 2D**). Finally, we investigated the influence of the definition of the interface on the results (**Fig. 2C**), by training and testing models on only the support and core (SC) interfacial residues, or only the core and rim (CR). The CR model yielded the best overall performance, and allowed to retrieve near-native conformations in the top 5 for almost all protein pairs (see also **Fig. S4**).

## 5 Discussion

In this work, we have investigated the possibility of evaluating complex candidate conformations by learning local 3D atomic arrangements at the interface. We have implemented a deep learning based approach that does not require rotational data augmentation, and that is sensitive to the relative orientations of the protein residues. We have shown that the method can help to improve the discrimination between near-native and incorrect conformations and could be useful in combination with more classical scoring functions. Our best results are based on a local representation of the interface and a distribution of patterns that the machine identifies at specific locations of the interface. The next tasks will be to more systematically explore the hyperparameter space, and to assess the performance of the method on a larger set of complexes. We will also address the issue of the multiple usage of protein surfaces. Indeed, two proteins may interact through different binding modes, depending on the context (presence of other chains for instance). The choice of one relevant binding mode, or the accounting for multiple binding modes is a difficult question. We envision many applications for the local-environment-based approach we propose, including the identification of physiological interfaces and the prediction of mutational outcomes on binding affinity.

## A Appendix

**Table S1:** PDB codes for the test set. The biological assembly ids are given in parenthesis. The extent of conformational rearrangement between the docked protein forms and the bound forms was assessed by the interface root-mean-square deviation (I-RMSD), computed on the C*α* atoms and after superimposition; none: the bound forms were used for docking, small: I-RMSD *≤* 1.5Å, medium: 1.5Å *<* I-RMSD *<* 2.2Å, and large: I-RMSD *≥* 2.2Å.

| Receptor | Ligand | Complex | Rearrangement |
| --- | --- | --- | --- |
| 1ku6_B | 1ku6_B | 1fsc_A:A (1) | medium |
| 1wsu_A | 1wsu_C | 1wsu_A:B (1) | small |
| 1li1_F | 1m3d_E | 1li1_F:E (1) | small |
| 1yy9_D | 1ck4_B | 2b2x_H:A (1) | large |
| 1li1_C | 1m3d_A | 1li1_C:B (1) | small |
| 1li1_C | 1m3d_B | 1li1_C:B (1) | small |
| 1m3d_H | 1m3d_H | 1m3d_H:G (2) | small |
| 1ezx_C | 2r9p_E | 2r9p_A:E (1) | medium |
| 1rkc_A | 1ydi_A | 2hsq_A:A (3) | large |
| 2c63_D | 2c74_A | 2c63_D:C (2) | small |
| 2c63_A | 2c74_A | 2c63_A:B (1) | small |
| 2vp7_A | 2vp7_A | 2yyr_A:B (1) | medium |
| 1m3d_G | 1m3d_H | 1m3d_G:H (2) | none |
| 1m3d_G | 1m3d_L | 1m3d_G:L (2) | none |
| 1li1_A | 1m3d_D | 1li1_A:B (1) | small |
| 2c9w_A | 2jz3_C | 2c9w_A:C (1) | small |
| 2z3q_B | 2z3q_C | 2z3q_B:A (1) | small |
| 1m3d_D | 1m3d_F | 1m3d_D:F (1) | none |
| 1i7x_B | 1i7x_C | 1i7x_B:C (1) | none |
| 3cwb_F | 3cwb_P | 3cwb_F:C (1) | small |

**Figure S1:**
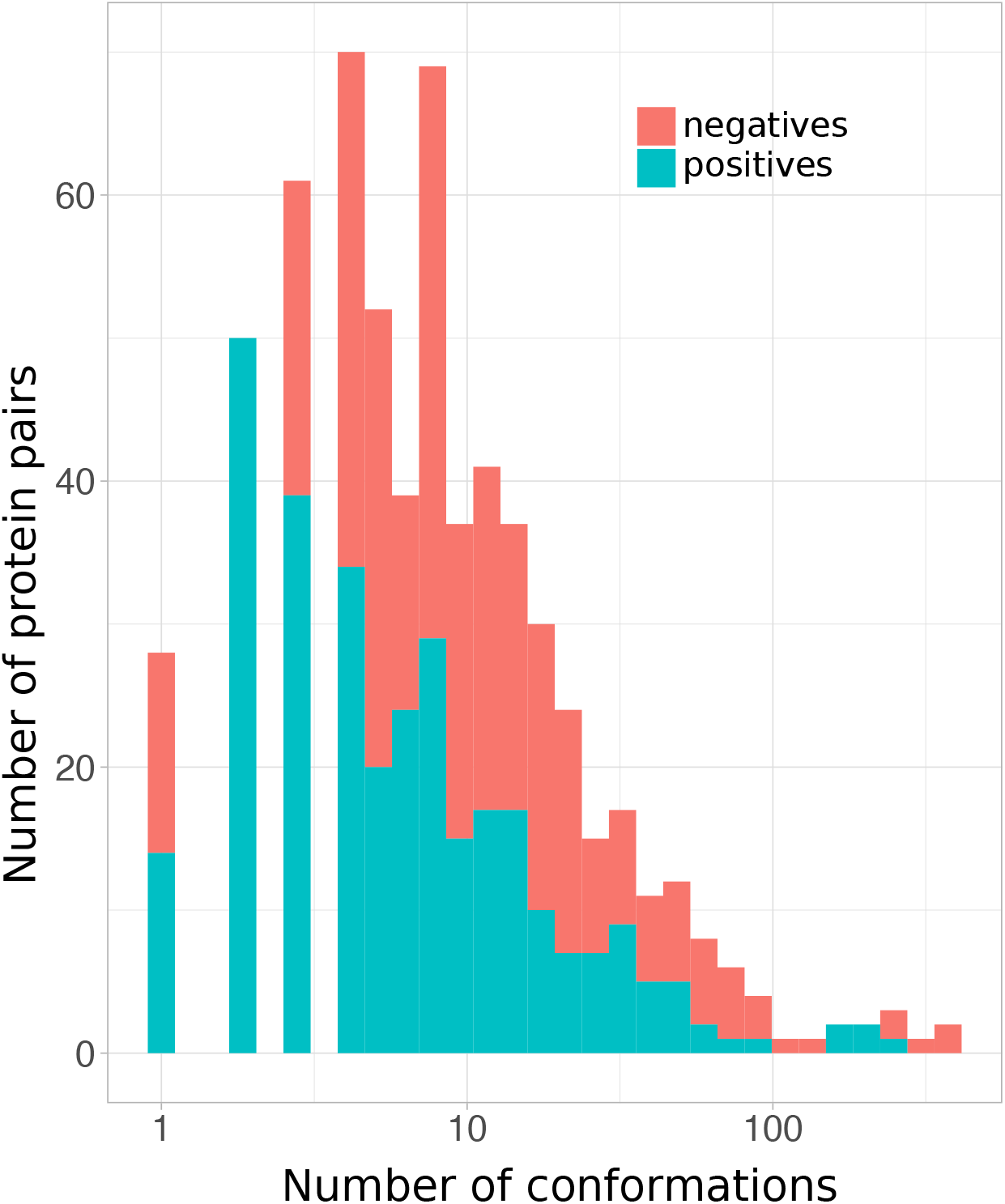
Cumulative distributions of positives (acceptable or better conformations) and negatives (incorrect conformations) in the training set.

**Figure S2:**
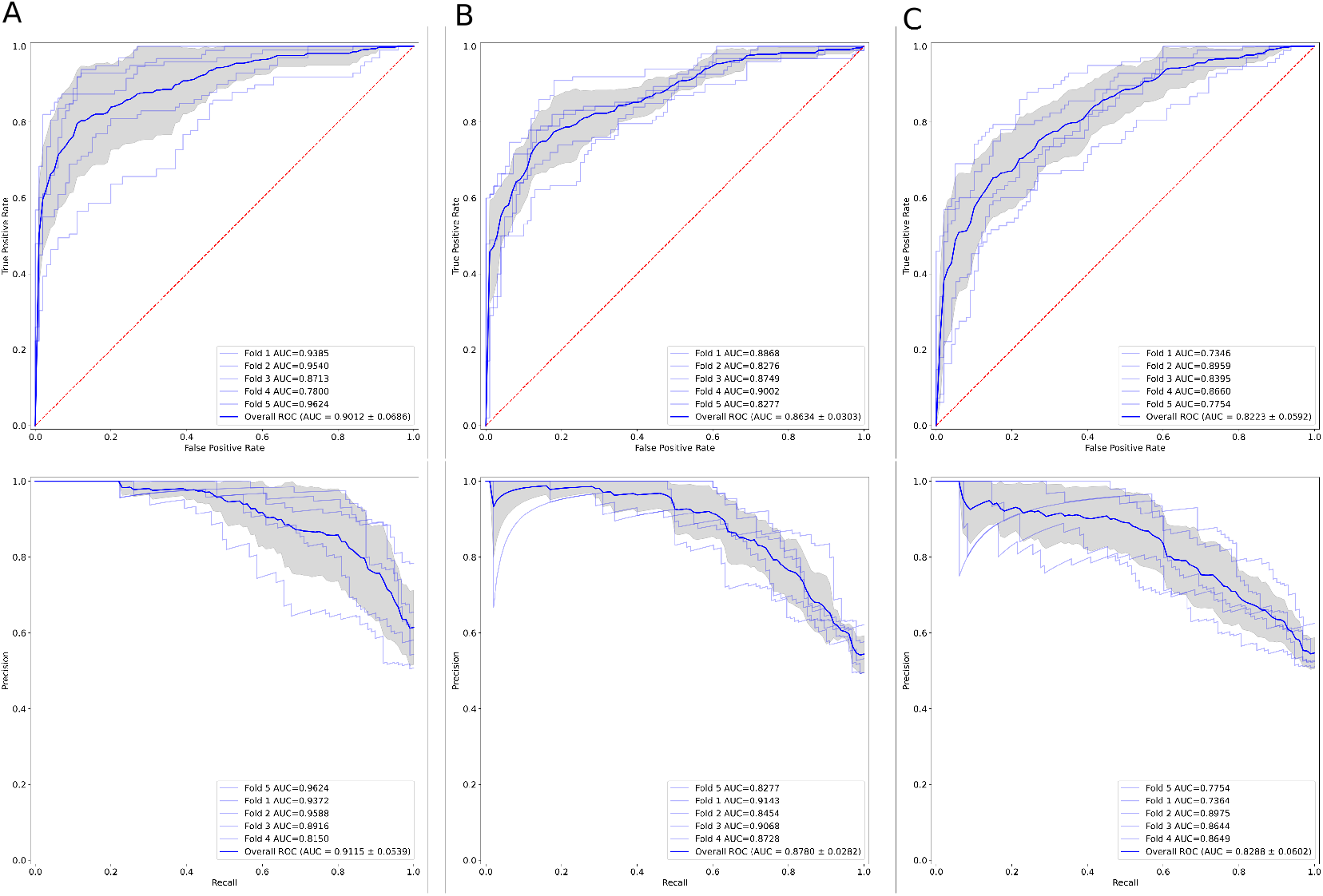
Validation ROC (on top) and PR (at the bottom) curves for 5 *x* 3 DLA-Ranker models. The models were trained and validated on entire interfaces (A), on the subset of core and rim interfacial residues (B), or on the subset of support and core interfacial residues (C).

**Figure S3:**
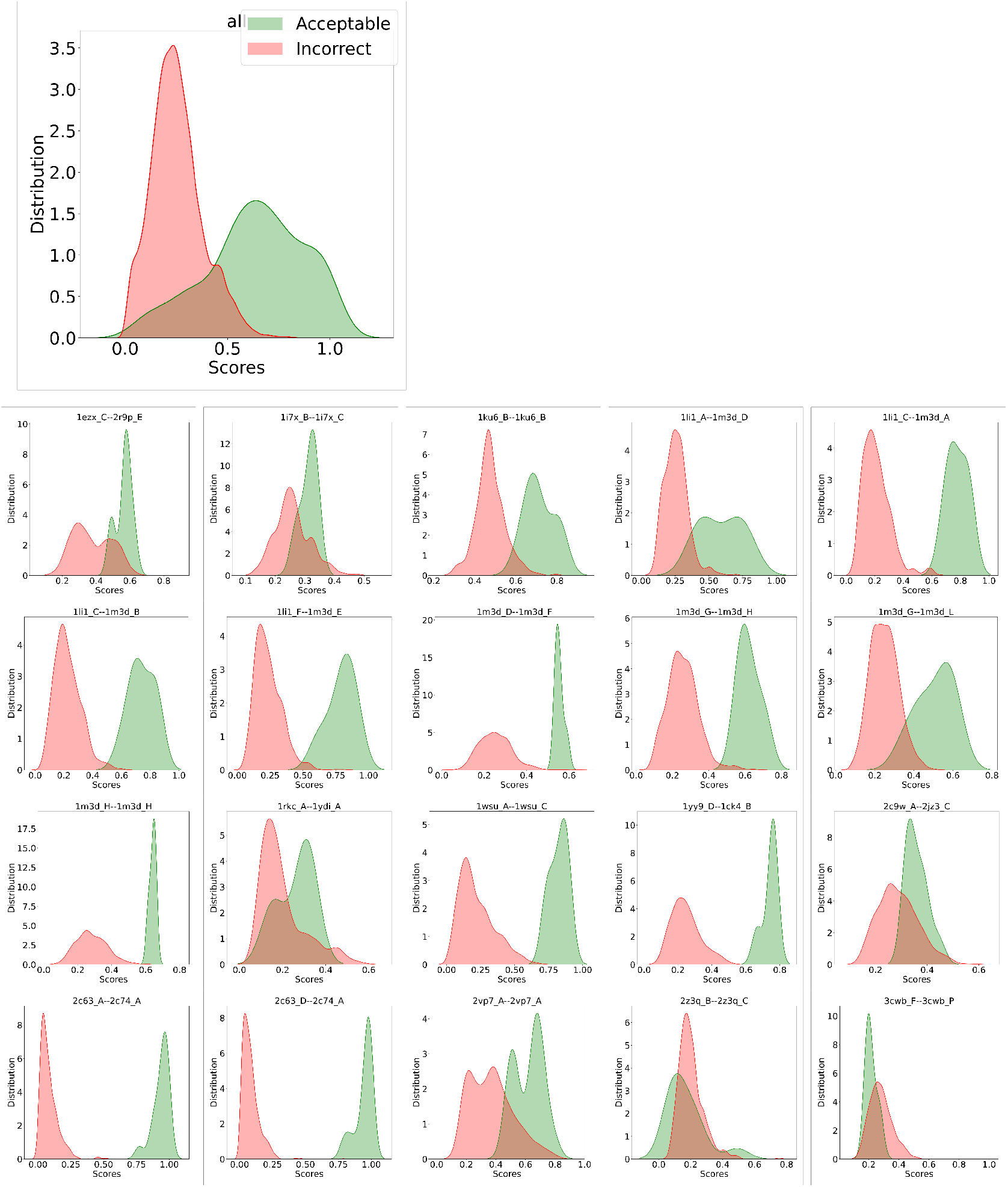
Distributions of DLA-Ranker *Q* scores computed for the test set conformations. We report the scores computed by one of the 5 DLA-Ranker trained models. The overall distributions are shown on top, and the per-protein-pair distributions below.

**Figure S4:**
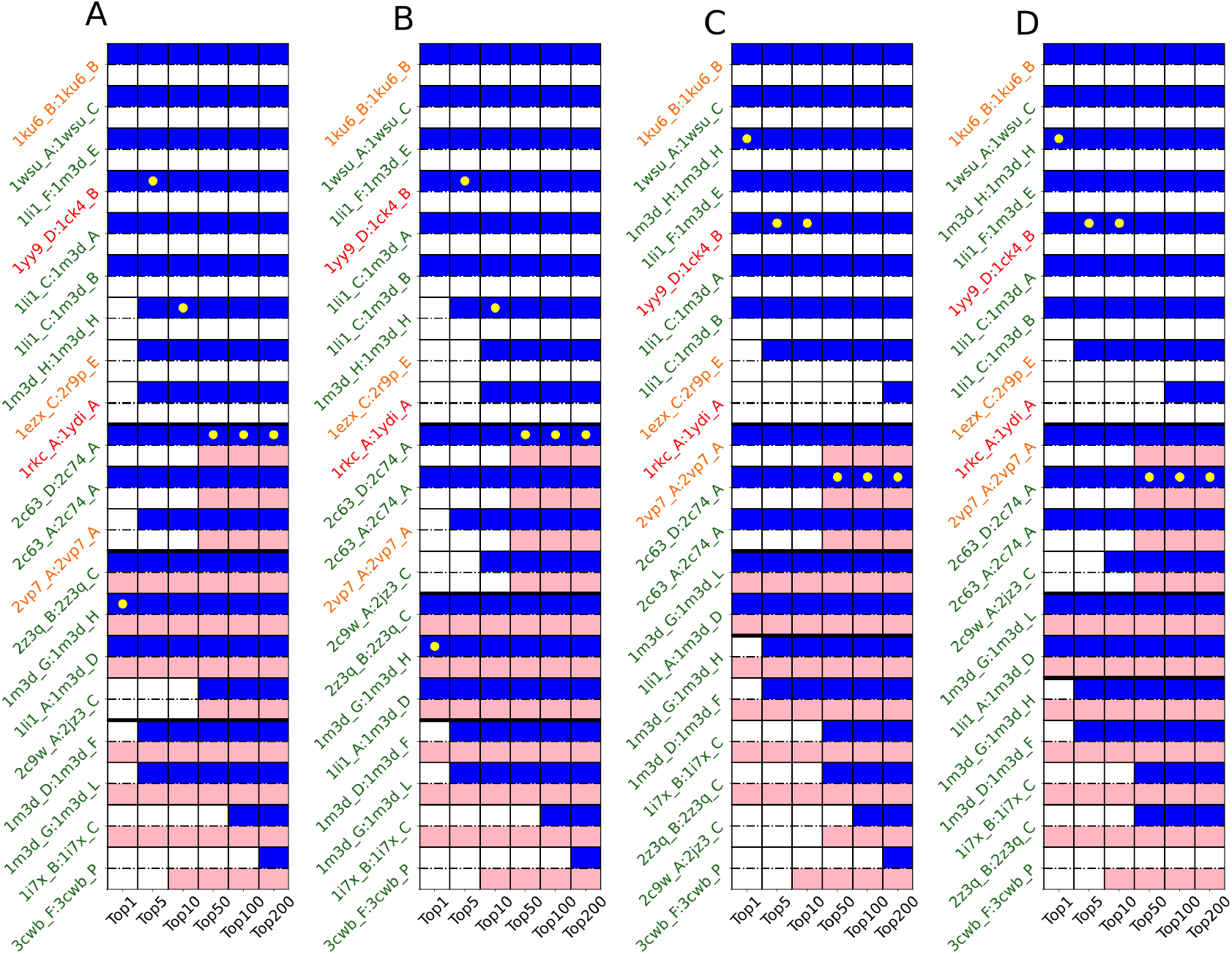
Ranking results per protein pair. **A**. DLA-Ranker trained and validated on core and rim residues only (DLA-Ranker-CR), **B**. DLA-Ranker-CR combined with CIPS, **C**. DLA-Ranker trained and validated on support and core residues only (DLA-Ranker-SC), **D**. DLA-Ranker-SC combined with CIPS. The color codes are the same as in Fig. 1.

